# Fluorescence polarization control for on-off switching of single molecules at cryogenic temperatures

**DOI:** 10.1101/204776

**Authors:** Christiaan Hulleman, Max Huisman, Robert Moerland, David Grunwald, Sjoerd Stallinga, Bernd Rieger

## Abstract

Light microscopy allowing sub-diffraction limited resolution has been among the fastest developing techniques at the interface of biology, chemistry and physics. Intriguingly no theoretical limit exists on how far the underlying measurement uncertainty can be lowered. In particular data fusion of large amounts of images can reduce the measurement error to match the resolution of structural methods like cryo-electron microscopy. Fluorescence, although reliant on a reporter molecule and therefore not the first choice to obtain ultra resolution structures, brings highly specific labeling of molecules in a large assemble to the table and inherently allows the detection of multiple colors, which enable the interrogation of multiple molecular species at the same time in the same sample. Here we discuss the problems to be solved in the coming years to aim for higher resolution and describe what polarization depletion of fluorescence at cryogenic temperatures can contribute for fluorescence imaging of biological samples like whole cells.

## 1. Introduction

Over the last decade new microscopy methods, frequently referred to as ‘super-resolution microscopy’ or ‘nanoscopy’, have achieved much higher resolution (~10-50 nm) than conventional light microscopy (~250 nm). The resolution of conventional microscopy is limited by diffraction to a length scale λ/2/NA, where λ is the emission wavelength, and NA=*n* sin(α) is the so-called numerical aperture of the objective, where *n* is the refractive index of the immersion medium and α is the marginal ray angle of the light beam collected by the objective. With the introduction of different nanoscopy techniques^[1–4]^ the diffraction limit has been circumvented, giving a resolution that starts to close the resolution gap between light and electron microscopy. Where electron microscopy can reveal details below the order of a nanometer it does not offer efficient specific labeling or multi-color imaging the way light microscopy does.

A group of techniques termed single-molecule localization microscopy (SMLM), e.g. PALM, (d)STORM, and many other flavors^[2–5]^ has found the most widespread use as it is relatively undemanding on the experimental setup and the subsequent image analysis. The key idea is to image and localize isolated single fluorescent molecules over many acquisitions. Essential is the ability to switch the fluorescent labels between a bright emitting “on”-state and a dark non-emitting “off”-state resulting in blinking. This leads to a sparse subset of all labels in the sample being in their “on” state at any time point which can be localized with a much higher precision than the diffraction limit.^[6]^ This localization uncertainty is on the order of λ/NA/√N_ph_, implying that already a moderate photon count of 100-1000 photons results in a ten times smaller uncertainty compared to the diffraction limit. The required ratio of on/off times to image only single emitters in a region of size λ/NA is typically smaller than 1/100 to 1/1000 depending on labeling density, exposure time and additional parameters.^[7]^

The image resolution, defined as the size of the smallest detail that can be reliably discerned in an image, is determined by the localization precision and the density of fluorescent labels. Previously we have introduced the concept of Fourier Ring Correlation (FRC) into super-resolution microscopy for taking all resolution factors into account.^[8]^ The FRC quantifies the available image information as a function of spatial frequency, i.e. across all length scales, and points to a resolution limit via a threshold criterion. The FRC resolution in SMLM can be a value of 2π times the localization uncertainty, depending on an adequate labelling density,^[8]^ providing a resolution of down to ~30 nm for high photon-yield fluorophores like Alexa647 (several thousand photons, localization precision ~5 nm).

There are a number of emerging developments that address the limitations of labelling technologies. Advances in bio-photochemistry have resulted in the development of labels that make a direct covalent bond to the target molecule, such as click-chemistry,^[9,10]^ effectively giving a small (<1 nm) Label size. Improvements of known dyes like Rhodamines has led to much brighter, more stable and cell permeable labels with tuneable emission properties.^[11,12]^ Also, techniques for high labelling density^[13]^ and the use of data fusion techniques^[14,15]^ ameliorate the labelling limitations. Based upon these developments, it may be anticipated that the localization imprecision will soon become the limiting factor for resolution.

Very recently, a new breakthrough technique to further push localization imprecision was proposed, called MINFLUX.^[16]^ In this technique a single fluorophore is illuminated by a doughnut beam a number of times (typically 4 times), where the position of the doughnut beam is changed from illumination to illumination across a region of size *L*. The number of photons per illumination is collected at a photo-diode and the set of photon counts is used in a triangulation procedure to estimate the position of the molecule. Surprisingly, the localization precision scales as *L*/√N_ph_, independent of the diffraction length λ/NA. By choosing *L* on the order of 50 nm an order of magnitude in precision can be gained compared to standard SMLM. The big drawback of this technique, however, is that the position of the molecule must be established to be within the tiny Region Of Interest (ROI) of size *L*~50 nm by a prior experiment. A development road to application of MINFLUX across the full FOV of the objective does not appear to be in sight, and seems less than straightforward to devise.

The localization precision can be reduced to well below 1 nm if more than 10^6^ photons are recorded. Collection of these amounts of photons is possible for uncaging dyes^[17]^ which today have been used infrequently, in DNA PAINT approaches where an effectively endless reservoir of dyes is imaged^[18]^, or through imaging at cryogenic temperatures.^[19–21]^ At cryogenic temperatures dyes have a very low rate of photobleaching and localization precisions below 1 nm are typical. For fluorescent proteins, however, freezing only offers a moderate increase in total photon count before bleaching.^[22]^ The increased localization precision is especially beneficial for data fusion techniques that rely on localizations to register or align different identical particles to obtain a super-super-resolved particle.^[15]^

A challenge to data fusion methods lies in the need for the sample to be immobilized during acquisition of individual images, but working with cells at cryogenic temperatures has the inherent advantage of fixation while the method of vitrification by plunge freezing is considered a very fast and mild fixation technique, compared to chemical fixation at room temperature^[23]^. With vitrification being used for cryo-EM sample preparation it also allows easier integration into correlative studies.^[23–25]^

On the downside, traditional switching mechanism for photon emission of dyes do not work for frozen samples. Photo-switching either requires a conformational change of the fluorophore itself such as in PALM^[2]^ or a liquid buffer solution for chemical interaction with the fluorophores such as in dSTORM.^[4]^ Very inefficient photo-conversion at cryogenic temperatures has been reported and used for correlative studies in TEM,^[22]^ but imaging of a densely labeled sample would not have been possible at such photoswitching rates. For a very small number of dyes spontaneous blinking has successfully been used to reconstruct a 3D structure from 2D projections.^[20]^

## 2. Fluorescence polarization control

Here, we propose the use of polarization control to establish on/off switching of fluorophores at cryogenic temperatures. At cryogenic temperatures, already in the range of liquid nitrogen, the absorption dipole axis of an individual fluorophore has a fixed orientation making it possible to tune the excitation of the fluorophores by rotating the state of polarization of the excitation beam (see Figure 1). The excitation efficiency scales as cos^2^θ with θ being theangle between the (linear) excitation polarization and the dipole axis. This effect alone gives rise to an on/off ratio of one to one, which however, is far from what is needed for producing the sparsity that is typically used in SMLM. The level of sparsity can be improved by making use of fluorescence depletion by stimulated emission, similar to STimulated Emission Depletion (STED).^[1]^ A second beam, red-shifted compared to the excitation beam, and with a polarization orthogonal to the excitation beam is used to illuminate the sample (see Figure 1). As a result most excited molecules are driven back to the ground state with the exception of molecules with an absorption dipole that is close to the orientation perpendicular to the polarization of the depletion beam. As a consequence, the fluorescence emission will no longer follow the original cos^2^α distribution but a much more sharply peaked distribution instead.

**Figure 1: a).**
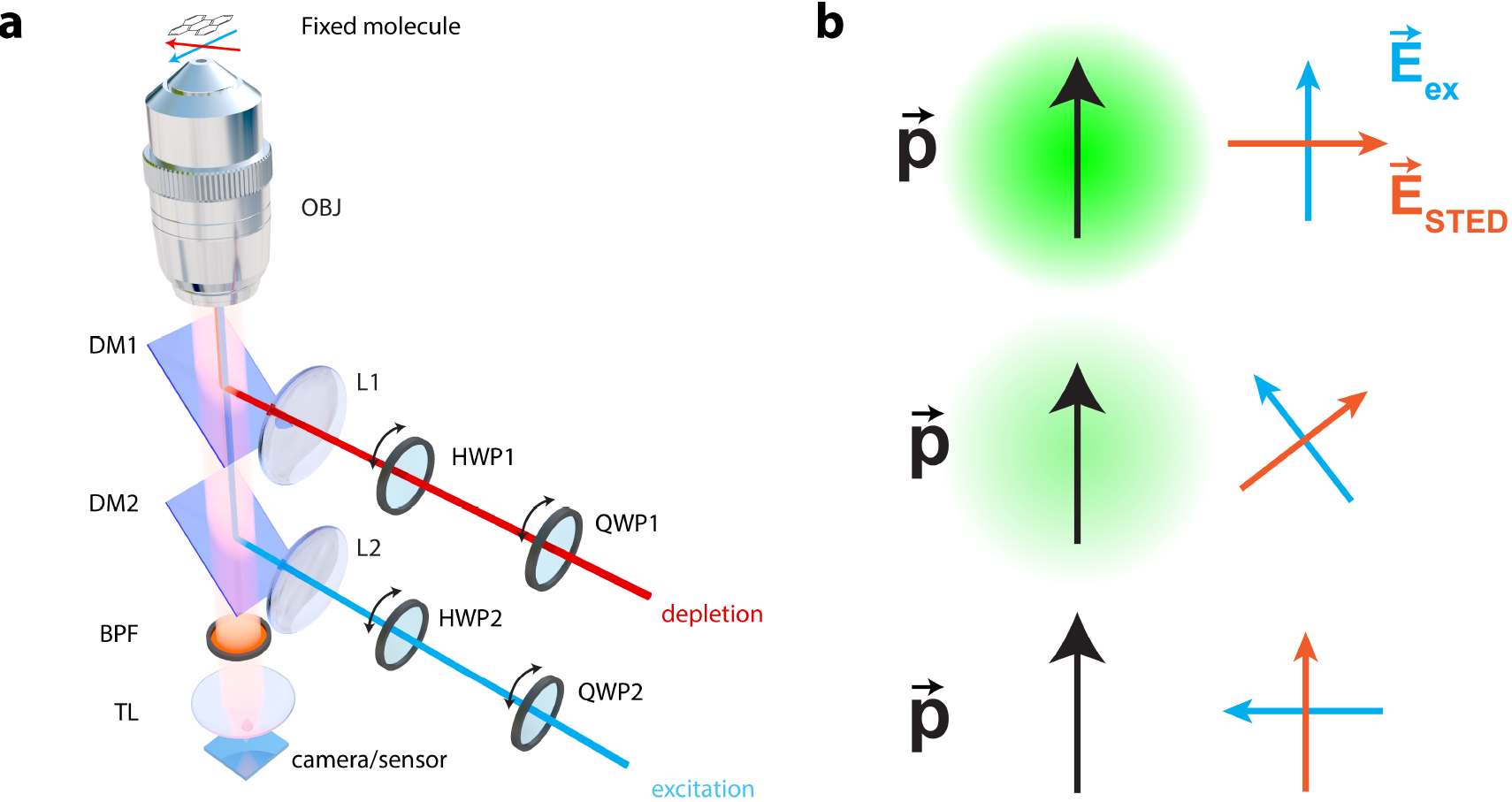
Schematic drawing of the setup. A custom built microscope is comprised of objective OBJ and tube lens TL. Dichroic mirrors DM1 and DM2 are used to reflect excitation and depletion laser light towards the objective, and let the fluorescence from the sample pass to the camera. A bandpass filter BPF is used to further suppress the potential leaked through laser light. Lenses L1 and L2 are used to focus the lasers on the back focal plane of the objective for epi-illumination. Full control over the polarizations of both lasers in the sample plane is obtained through the use of half-lambda (HWP1 & HWP2) and quarter-lambda wave plates (QWP1 & QWP2) for each laser path, respectively. **b)** With the excitation laser 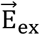 parallel to the molecular transition dipole 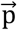 and the depletion laser 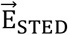 perpendicular, the molecule fluoresces at maximum intensity. The excitation efficiency is reduced when 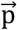 is at an angle with 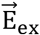. Moreover, the increased inner product with 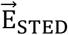 leads to efficient depletion of the molecule and a sparsity in the sample based on angular selection of transition dipoles, compared to a case where only 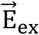 is present.

We can make an estimate of the improvement in sparsity by employing the analogy of the proposed method with STED. Hell^[26]^ stated that the resolution of STED scales as 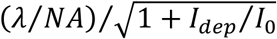, where *I*_*dep*_ is the intensity of the depletion beam and *I*_0_ ≈ ℏω/*στ* (with ℏω the photon energy, *σ* the crosssection, and *τ* the lifetime) a constant intensity. Similarly, we conjecture that the angular emission profile after depletion has a Full Width Half Maximum (FWHM) that may be described by: 
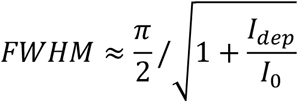
resulting in a sparsity factor π/FWHM that can be much larger than 1, in principle achieving the same type of FWHM narrowing as in standard STED. So far, we have only considered excitation and depletion polarizations and dipole orientations in the plane of the sample. This matches with the relatively low NA of objective lenses that can be used in cryogenic setups (typically up to NA=0.7).^[19,21]^ Under these conditions the polarization of the excitation and depletion beams are necessarily close to in-plane polarized, and fluorophores with dipole orientations that are substantially tilted with respect to the sample plane are not excited and remain invisible. Possible extensions to high-NA excitation, depletion, and imaging using immersion technology, in particular a Solid Immersion Lens (SIL), would open up possibilities for polarization control over the full 4π solid angle of polarization and dipole orientations.

Hafi et al. already put forward the idea to use polarization as an effective way to introduce sparsity in combination with STED, similar to our setup in Figure 1.^[27]^ Later Frahm and Keller showed however that their results were in fact not based on polarization control but due to post processing of the data by a sparsity enhancing deconvolution algorithm.^[28]^ Closer inspection of the data of Hafi and co-workers reveals that they reported very poor modulation depths when changing the polarization (their Figure 2). Two experimental factors might have caused these poor modulation depths, thereby impairing their results, while the conceptual idea is working, as we demonstrate below. To do so we first improved the effective polarization in the sample plane (the first experimental factor), by carefully calibrating the different optical elements in the microscope such that full modulation over background can be seen. Second, we used fluorescent molecules fixed by spincoating in a polymer solution to have stable molecular transition dipoles (the second experimental factor).

**Figure 2:**
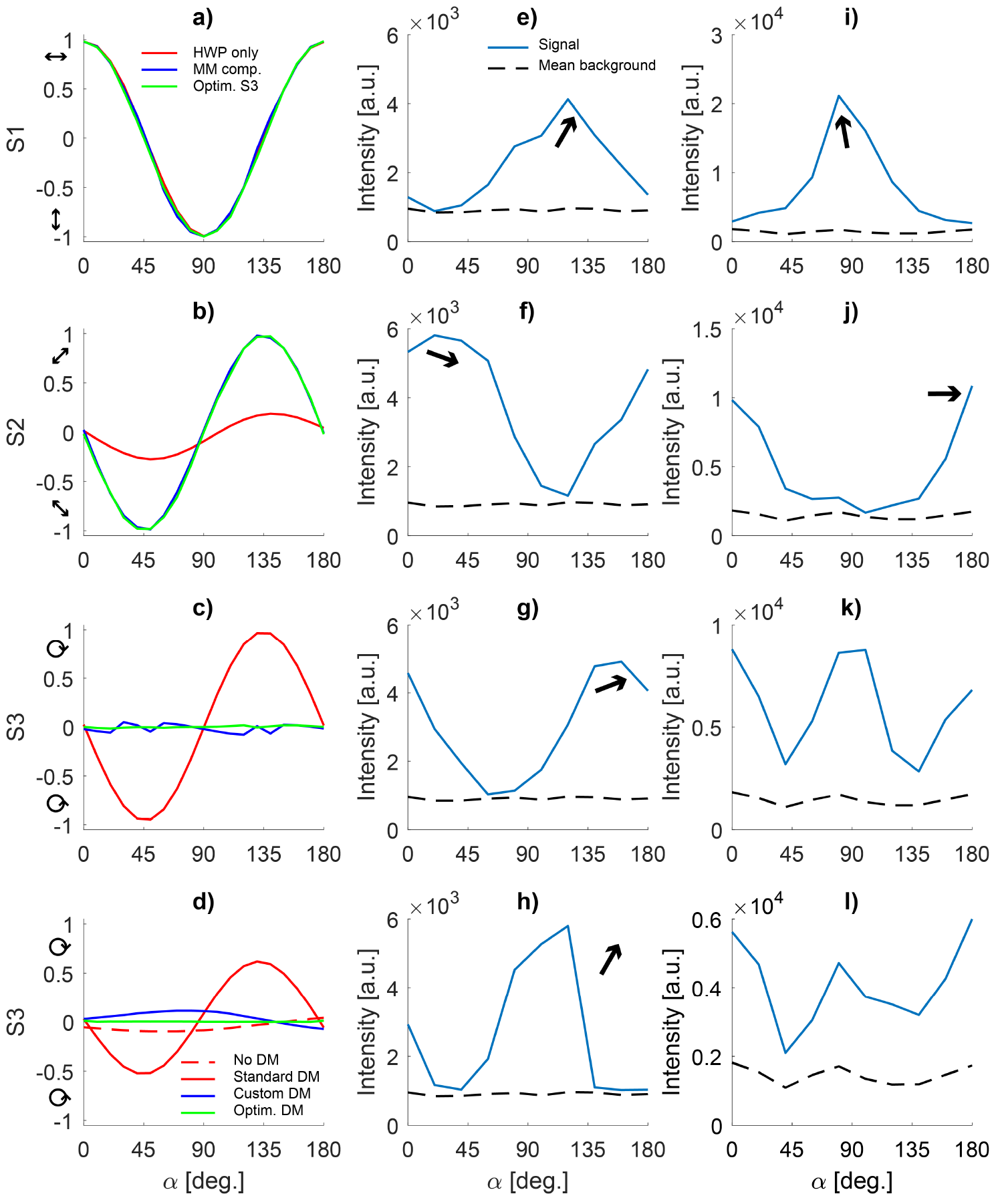
**a, b, c)** Normalized Stokes vector components S1, S2 and S3 for the dichroic mirror used for excitation as a function of α the desired polarization orientation angle. Rotating only a HWP yields the red curve. Using Calculated orientations of the QWP and HWP based on the measured Mueller matrix yields the blue curve. Optimizing the orientation of the WP based on the resulting Stokes vector yields the green curve. **d)** Normalized S3 Stokes vector component for the standard and custom dichroic mirror. The dashed red curve is a direct measurement of a rotating HWP only. **e, f, g, h)** Mean intensity of single molecule emitters using the optimized QWP and HWP orientations, the black arrows depict the estimated molecule's transition orientation. **i, j, k, l)** Mean intensity fluctuation of single molecule emitters during rotation of only a HWP, again the black arrows indicate estimated molecular transition dipole orientations.

### 2.1 Optimizing light polarization at the sample

We developed a method to control the polarization of excitation and STED beam at the sample plane. To this end we calibrate the optical system to know how polarization is changed by the different components in the setup as depicted in Figure 1, and in particular we measured the effect of the retarders (HWP and QWP) instead of assuming their nominal action. In the following we shortly introduce the concept of Stokes vector and Mueller matrix, which allows the description (and manipulation) of the polarization state of light. We describe the electric field of a monochromatic plane wave, traveling along the z-axis, by: 
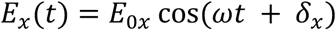

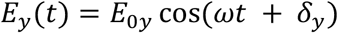

Here we have taken z = 0 for convenience and *E*_0x_ and *E*_0y_ represent the amplitude of the electric field in the *x*- and *y*-direction, respectively. The angular frequency is given by ω, and δ_x_ and δ_y_ are the phase factors for the respective electric field components. In general, the tip of the electric field vector traces an ellipse in space for arbitrary amplitudes and phases, the polarization ellipse. It can be shown that the Stokes 4-vector *S* and the electric fields are related as:^[30]^ 
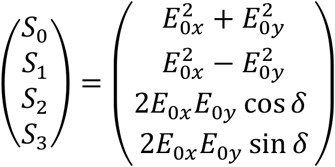
with *δ* = *δ*_*y*_ − *δ*_*x*_. The first stokes parameter *S*_0_ describes the total intensity of the light. Of the remaining parameters, *S*_1_ describes the amount of horizontally or vertically polarized light, *S*_2_ the amount of diagonally polarized light (±45°) and *S*_3_ the amount of left/right-handed circularly polarized light. The 4×4 Mueller matrix **M** transforms two Stokes vectors *S*_out_= **M** *S*_in_ and describes how the polarization state is changed by an optical system. Once the Mueller matrix of a system has been determined, the required input polarization state in order to get a desired output polarization state can be retrieved via **M**^−1^. Here, we have used a dual rotating retarder polarimeter,^[29]^ which consists of a polarizer, a QWP, the system under test, a second QWP and an analyzer. The two QWPs are rotated simultaneously but with a 5:1 ratio which gives the 16 elements of M after Fourier analysis of the resulting intensity signal for different angles. With the free space requirement that **M** is the identity matrix we can calibrate the polarimeter^[31]^ and the orientation and retardance of the retarders. In our implementation, the QWPs (AQWP05M-600, Thorlabs) are rotated with computer-controlled rotation stages (8MPR16-1, Standa). The polarizers are of the GL10 type (Thorlabs) with an anti-reflection coating suitable for the wavelength used.

In order to measure *S*_in_ and *S*_out_, we use a Stokes polarimeter^[30]^ comprised of a rotating QWP (AQWP05M-600, Thorlabs) and a polarizer (GL10, Thorlabs). Here, the retarding wave plate is rotated with a computer-controlled stage (8MPR16-1, Standa) as well. The laser light running through the system is modulated with the aid of an optical chopper wheel (MC2000-EC, Thorlabs) and detected with a silicon photo diode (DET10A, Thorlabs), after focusing with an additional lens (LB1901-B, Thorlabs). A lock-in amplifier (SR830, Stanford Research Systems) is used to demodulate the detector signal. Both types of setup are computer-controlled from within MATLAB (The MathWorks). Thus, with the combination of a Stokes and a Mueller polarimeter, we can fully determine the polarization transfer function of our microscope (see Figure 1a) and the polarization state of the light itself in the sample plane.

### 2.2 Results of polarization control

In order to evaluate our calibration procedure over naive usage of the nominal action of the retarders, we measured Stokes vectors for both setups (see Figure 2a-c). Without DM1 and QWP2 present (see Fig 1a), we insert 638 nm linearly polarized laser light into the excitation path (LDH-D-C-640, PicoQuant), where the initial linear polarization is generated with a Glan-Laser calcite polarizer (GL10, Thorlabs). First, we naively try to rotate the polarization by rotating an achromatic half-wave retarder (AHWP05M-600, Thorlabs) in steps of 5° and measure the polarization state at the sample plane with Stokes vector polarimetry (compare the red solid line in Figure 2a-c). In the coordinate system of the microscope, a horizontal polarization, corresponding to *S*=(1,1,0,0) is identical to *s*-polarized light hitting the dichroic mirror DM2 (FF652-Di01, Semrock). This corresponds to α=0° in Figure 2. Vertically polarized light is identical to *S*=(1,−1,0,0) and corresponds to α=90°. From Figure 2a-c we see that these two linear polarizations are well-defined with normalized S=(1, 0.981, 0.013, 0.023) ± (0,2,1,1) × 10^−3^ (1σ) at α=0° and *S*=(1, −0.995, −0.093, −0.005) ± (0,0.3,2,2) ×10^−3^ at α=90°. However, for the diagonal polarizations at α=45° and α=135°, corresponding to *S*=(1,0,∓1,0), it is obvious that *S*_2_ is far from (minus) unity, and in fact *S*_3_ is near (minus) unity. Therefore, in the straight forward implementation, polarizations deviating from horizontal and vertical are increasingly elliptically polarized, even approaching circularity.

Next, we calibrate our Mueller polarimeter and measure the Mueller matrix of the excitation path with the method described elsewhere.^[31]^ In this case, we do not insert HWP2 and QWP2 into the path. Therefore, we obtain the Mueller matrix of the combination of the dichroic mirror DM2, lens L2 (LA1433-633, Thorlabs) and the objective (Plan Apo VC 100x/1.40, Nikon), as well as a measurement of the retardances of the used QWPs. Here we used QWP2 as part of the Mueller polarimeter, thus obtaining the actual retardance. Furthermore, we determine the retardance of HWP2 with the method described by Goldstein.^[30]^ The total polarization transfer function of the excitation light is then **M**_tot_=**M**_mic_ **M**_HWP2_ **M**_Qwp2_, where **M**_Qwp2_ and **M**_HWP2_ are the Mueller matrices of the wave plates, and **M**_mic_ is the Mueller matrix of DM2 with the objective. By this procedure we are able to predict the Stokes vector, and hence the polarization, of the light moving through the excitation branch including HWP2 and QWP2. Moreover, we can predict the angles of the retarders that are necessary to obtain any required polarization state in the sample plane. We use this possibility to compensate the error in polarization, induced by DM2 and the objective, in order to obtain linearly polarized light in the sample plane (compare the blue solid line in Figure 2a-c). From visual inspection, it is clear that over the whole range of linear polarizations, the ellipticity is strongly reduced (compare blue and red line in Figure 2c).

We focus our attention on the diagonal polarizations which are problematic without correction. For α=45°, S=(1, 0.015, −0.976, −0.015) ± (0, 2, 2, 2) ×10^−3^ and for α=135°, S=(1, 0.025, 0.989, −0.029) ± (0, 2, 3, 1) ×10^−3^, after calibration. The linearity of the polarization at these angles therefore is vastly improved. We see in Figure 2c) that there still is some residual ellipticity with the Mueller-matrix based compensation method. We therefore add a numerical optimization algorithm, which searches for the optimal WP angles to minimize *S*_3_. The cost function returns the error between the ideal and the measured Stokes vector components: err = |*S*_*1*_−*S*_*1m*_|+|*S*_*2*_−*S*_*2m*_|+10×|*S*_*3*_−*S*_*3m*_|, where we emphasize the error in *S*_*3*_, the component which leads to ellipticity. For each angle α, the optimization is run (compare the green solid lines in Figure 2a-c). Ideally, in all cases *S*_*3*_ is identical to zero. Visibly, *S*_*3*_ is closer to zero throughout the range of α after optimization. The root-mean-square error for the different schemes in Figure 2c) shows a substantial improvement from 0.66 (naïve rotation of HWP2 - red), 0.04 (Mueller matrix - blue) and 0.01 (optimized - green).

Figure 2d) shows the *S*_*3*_ component of a Stokes vector measurement of just a HWP (dashed red line). When compared to the case that the DM is installed (red solid line), the dramatic effect of the DM on the ellipticity of the polarization is clearly visible. We obtained an optimized DM from Chroma, not only for a high reflectivity of the laser, but also for a minimal phase difference between the *s*- and *p*-components, which indeed has nearly no influence on the ellipticity of the beam (compare measured blue curve), and merely changes the circular component, induced by the HWP, in handedness. An additional optimization step as described above essentially eliminates this part (green curve).

We now turn to single-molecule fluorescence measurements of fixed dyes (at room temperature) as a function of polarization to investigate the performance of our calibration above. We apply and compare the naive polarization rotation scheme and the calibrated, optimized scheme. For each angle α we obtain a fluorescence image with the aid of an EMCCD camera (Ixon Ultra 888, Andor), with an exposure time of 1 second, an EM gain of 25 and a sensor temperature of −60°C. In order to prepare the samples, a stock solution of ATTO 647N (ATTO-TEC) was diluted to a final concentration of 2.8×10^−10^M in demineralized water, mixed with 0.5 w% of PVA. Spin coating this solution at 3000 RPM for 1 minute onto cleaned cover slips resulted in samples of single molecules with a randomly fixed dipole transition moment. Figure 2e-h) show the mean fluorescence intensity of a selection of 4 molecules for the optimized polarization rotation scheme (blue solid line). The background (black dashed line) has been determined as the average of 5 locations without any molecules. The black arrows serve as an illustration of the in-plane dipole moment of each molecule, based on the phase of the intensity modulation. Figure 2i-l) show the results for the naive implementation on 4 other molecules. Here we clearly see the improvement of our calibration procedure. For (near) horizontal and vertical dipole orientations, the modulation depth is comparable in both cases (nearly 100%), as for these polarizations the DM hardly alters the polarization. However, for diagonal transition dipole moments, the calibration approach remains near a 100% modulation depth, highlighting the superiority of our scheme, in contrast to the naïve implementation where the near-circular quality of the polarization at the relevant angles α results in a strong electric field component along the dipole transition moment.

## 3. Cryostat design

Among the various methods to boost the number of available photons cooling of the sample to cryogenic temperatures, which also fixes the dipole moments of all emitters in the sample, is ideally suited for the use of polarization STED to enable high density labeling while maintaining emitter sparsity. However, use of a STED beam adds a heat load not typically present in imaging at cryogenic temperatures. A very powerful cryostat design, enabling cryoFM on practically any microscope platform, was presented by Li et al‥^[21]^ Adding an additional heat source to cryoFM raises the question what can be done to improve the cooling capacity of such a cryostat, both with respect to transferring heat from the sample and maintaining imaging duration upwards of five hours.

Different methods have been suggested to increase the cooling capacity to the sample. The modularity of the Li cryostat is of high practical value as it demands a minimum of modifications to the microscope platform, arguing to focus on increasing the tank volume, raising questions on how to best scale the size of the heat exchange interface in the cryostat (see Figure 1 in Li et al.) while maintaining the structural stability of the original design. We used finite-element simulations to optimize cooling capacity and mechanical stability. Simulations were done in Solidworks Simulation (Premium package, Dassault Systems) using a curvature-based solid mesh with 4 Jacobian points and element sizes between 1 mm and 20 mm. For thermal simulations about 433.000 nodes, forming roughly 270.000 elements, with a maximum aspect ratio of ~200 (>90% of elements had a ratio <3 and less than 0.5% of elements had a ration >10) were used. For frequency simulations, finding the first five natural frequencies and their amplitude at resonance, about 170.000 nodes, forming > 100.000 elements, with a maximum aspect ratio of ~48 (>89% of elements had a ratio <3 and less than 0.6% of elements had a ratio >10) were used. The goals for a cryostat with an increased cooling capacity (see Figure 3) are to better tolerate the use of additional external heat sources, like a STED beam, possibly increase the duration of the cooling cycle to allow for longer observation times, and maintain cooling temperatures suitable for cryoFM using liquid nitrogen while maintaining maximum vibration damping.

**Figure 3:**
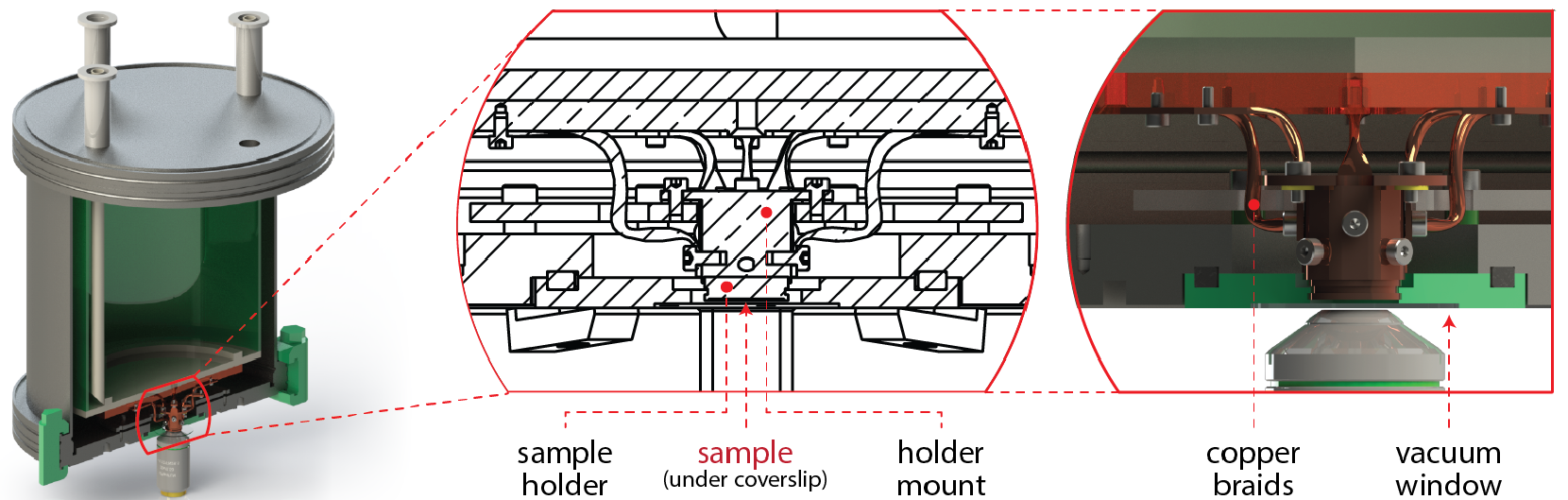
Improved cryostat design for cryo-fluorescence microscopy. The model depicted here features three main improvements over the design by Li et al. (Li2015): (1) a 2x larger liquid nitrogen tank for long-term super-resolution imaging, (2) four additional copper braids between the sample holder sample holder mount and the heat dissipater to allow more heat flux and increase cooling capacity and (3) vibrational mode optimization to maximize damping and minimize drift while maintaining thermal insulation. These improvements make the new design more suitable for cryo-polarizationSTED imaging.

By increasing the number of vibrationally decoupling thermal conductivity braids between the sample holder and the heat dissipater at the bottom of the nitrogen tank (Figure 3), the maximum heat flux is effectively doubled. We increased the size of the tank by a factor 2, allowing approximately double the original cold measurement time at traditional operation. The dimensions of the parts that suspend the sample holder in vacuum were optimized to increase thermal insulation and effectively dampen low-frequency oscillations, pushing the predicted resonance modes well into the kHz range. Specifically, we analyzed the impact of additional mounting points for the sample holder. We find that depending on the number of spacer rings an increase of mounting holes from 4 to 8 will result in a 2 K temperature increase at the sample holder while causing 32 K higher temperatures at the cooling disk. As to be expected, a higher number of mounting points yields higher resonance frequencies. Differences in the first three modes (along the *x*, *y* and *z* direction) were <30% with resonance frequencies in the range of 1900 Hz (*x*, *y*) and 2600 Hz (*z*) if 8 mounting points were used and 1400 Hz (*x*, *y*) and 1900 Hz (*z*) if 4 mounting points were used. The 4^th^ and 5^th^ modes showed differences below 1% and were in the range of 2500 Hz. Interestingly, the shape of the spacers used (strips vs. rings) was shown to be in favor of rings (yielding >20% differences in heat conductivity, roughly scaling with surface area differences). Taken together, optimization of cooling capacity of a closed cryostat as described by Li et al.^[21]^ should use fewer mounting holes as the stability improvements lie in a frequency range well above 1 kHz, but increased numbers of mounting holes increase heat flow substantially. While the effective heat load on the cryostat introduced by a STED beam is currently unknown our simulations indicate no downside to increasing the thermal coupling of the sample holder by increasing the number of copper braids between sample holder sample holder mount and the heat dissipater.

While cooling performance and the resulting imaging time remain to be assessed for STED experiments, this design should be capable of dissipating a substantial part - if not all - of the heat added by a depletion beam. Sensors that accurately register the temperature cooling stages within the cryostat can be used to examine the performance during experiments, as well as to provide an estimate of the amount of energy deposited into the sample by an external source.

## 4. Outlook & Conclusion

Minimizing the localization uncertainty and reducing the overall measurement error could lead to the FRC resolution values below or in the range of the size of an individual emitter. While new measurement modalities like MINFLUX show the way to improve localization uncertainty at low photon count, increasing the photon budget by means of improved labels or cooling of the sample are already possible today. The power of data fusion to reduce the measurement error has been demonstrated by resolving structural details of nuclear pore complexes in cells using SMLM style imaging.^[14]^ We showed that by using Stokes and Mueller polarimetry we were able to measure and improve the effective polarization in the sample plane. We considered design limitations to increase heat flow and cooling capacity of a modular cryostat to account for the additional heat load of a STED beam in cryoFM and argue that our measurements of the effective polarization depletion in the sample plane is the missing element to achieve nanometer ranged FRC resolution in SMLM.

Given the biological tools to unequivocally label specific proteins inside cells using CRISPR technology and to target different RNA populations very efficiently, we predict that SMLM imaging with resolutions corresponding to the size of individual emitters can provide a new level of understanding of the workings of the cell, for instance in gene expression and nuclear structure. For example in the field of chromosome capture where the correlation between gene location within the nucleus and expression is studied, currently only indirect tools exist to pinpoint differences between a cell in steady state and a cell that is activated. With our proposed setup it should be possible to label the genomic location of a gene (using for instance CRISPRainbow^[32]^ and then map that location in nm resolution during times of steady state and after activation. New labeling techniques like CRISPRainbow currently can target about 1000 specific genomic loci with sufficient repeat numbers. Using color barcoding as many as 6 loci can be visualized simultaneously. Single color SMLM imaging with nanometer range resolution would allow to use the number of guide RNAs targeted at a specific locus to distinguish different loci and boost the number of genomic substructures that can be resolved simultaneously by at least an order of magnitude. Similarly, number based barcoding with single guide RNA resolution would give us the ability to resolve the spatial relation between TADs and LADs at the level of individual genes.^[33]^ Ultimately, being able to locate specific genomic loci using CRISPR based labeling one could envision how data fusion can be used to resolve DNA structure based on resolving individual DNA intercalating dyes. While multi-color cryoFM with nanometer precise image registration has not been shown yet, it is easy to envision another class of experiments that would be enabled with it.

Another example, in the field of gene expression, it is still unclear how many copies of each export factor is loaded on a single mRNA allowing for it to be exported from the nucleus to the cytoplasm for expression. It is unclear what the threshold of factor loading is, and if there is a correlation between mRNA localization within the cell and factor dissociation, or if this is simply a factor of time. DBP5 is a DEAD box helicase, for which we have a deep understanding of its function from translation to mRNA nuclear export.^[34,35]^ Fundamental aspects of the relationship between mRNAs and DBP5, however, such as when and how many copies are associated and how this number varies between mRNAs - are poorly understood.^[36]^ Multicolor SMLM imaging with nanometer resolution will allow us to understand the occupation of nuclear pore complexes with DBP5, and other transport related molecules, and the loading state of DBP5, including the copy number as a function of subcellular localization of the mRNA, at a level currently out of reach. DBP5 is one example, but although many factors involved in mRNA export have been identified, we still have limited knowledge of the structural composition of mRNPs and its functions.^[37]^

## Acknowledgements

We thank Weixing Li and Jörg Enderlein for helpful discussion. B.R., C.H. and R.M. acknowledge European Research Council grant no. 648580 and B.R., M.H. and D.G. acknowledges National Institute of Health grant no. U01EB021238.

